# Why FIT and bHLH Ib interdependently regulate Fe-uptake

**DOI:** 10.1101/2022.02.12.480172

**Authors:** Yuerong Cai, Yujie Yang, Huaqian Ping, Chengkai Lu, Rihua Lei, Yang Li, Gang Liang

## Abstract

FIT (FER-LIKE IRON DEFICIENCY- INDUCED TRANSCRIPTION FACTOR) and four bHLH Ib transcription factors (TFs) bHLH38, bHLH39, bHLH100 and bHLH101, are the master regulators of Fe uptake genes, and they interact with each other to activate the Fe uptake systems. However, it remains unclear why FIT and bHLH Ib depend on each other to regulate the Fe deficiency response. By analyzing Fe deficiency phenotypes and Fe uptake genes, we found that the quadruple *bhlh4x* mutants (*bhlh38 bhlh39 bhlh100 bhlh101*) mimic the *fit* mutant. Subcellular localization analyses indicate that bHLH38 and bHLH39 are preferentially expressed in the cytoplasm whereas bHLH100 and bHLH101 in the nucleus. Transcriptome data show that the genes involved in Fe signaling pathway show the same expression trends in *bhlh4x* and *fit*. Genetic analyses suggest that FIT and bHLH Ib depend each other to regulate the Fe deficiency response. Further biochemical assays indicate that bHLH Ib TFs possess the DNA binding ability and FIT has the transcription activation ability. This work concludes that FIT and bHLH Ib form a functional transcription complex in which bHLH Ib is responsible for target recognition and FIT for transcription activation, explaining why FIT and bHLH Ib interdependently regulate Fe uptake.

## Introduction

Fe is one of the micronutrients crucial for plant growth and development because Fe acts as a cofactor involved in chlorophyll biosynthesis, photosynthesis, respiration, and other biochemical reactions. Fe deficiency often leads to Fe-deficiency symptoms such as interveinal chlorosis in leaves and reduction of crop yields. Plants absorb Fe from soil; however, Fe acquisition is challenging due to the low solubility of Fe in soil solution (Guerinot and Yi, 1994).

The requirement for efficient acquisition of Fe from soil has led to the evolution of two distinct uptake strategies, strategy I in non-graminaceous plants and strategy II in graminaceous plants (Marschner & Romheld, 1986; Romheld & Marschner, 1986; Grillet & Schmidt, 2019). Arabidopsis plants employ the strategy I which involves three sequential steps, mobilization of ferric Fe, reduction of ferric Fe, and transport of ferrous Fe. Insoluble ferric Fe is mobilized by acidification of the rhizosphere resulting from protons secreted by AHA2, a proton ATPase (Santi and Schmidt, 2009) and by phenolic compounds (Rodríguez-Celm et al., 2013; Schmid et al., 2014; Fourcroy et al., 2016; Tsai et al., 2018). Ferric Fe is then reduced to ferrous Fe by FRO2 (FERRIC REDUCTION OXIDASE 2) (Robinson et al., 1999), and finally translocated into roots by epidermis localized IRT1 (IRON-REGULATED TRANSPORTER 1) (Varotto et al., 2002; Vert et al., 2002).

FIT is a key regulator of strategy I since its loss-of-function causes reduction of Fe uptake genes including *IRT1* and *FRO2* and severe Fe deficiency symptoms (Colangelo & Guerinot, 2004; Jakoby et al., 2004; Yuan et al., 2005; Schwarz & Bauer, 2020). Interestingly, early reports showed that the singular overexpression of *FIT* does not result in constitutive activation of *IRT1* and *FRO2* (Colangelo & Guerinot, 2004; Jakoby et al., 2004). Later studies revealed that FIT interacts with each of four bHLH Ib transcription factors (TFs), bHLH38/39/100/101, and dual overexpression of bHLH Ib and *FIT* constitutively activates *IRT1* and *FRO2* (Yuan et al., 2008; Wang et al., 2013), implying that FIT and bHLH Ib TFs function synergistically. Unlike singular *FIT* overexpression, singular overexpression of *bHLH39* (or *bHLH101*) activates the expression of *IRT1* and *FRO2* (Yuan et al., 2008; Wang et al., 2013). However, Naranjo-Arcos et al. (2017) revealed that the activation of *IRT1* and *FRO2* by *bHLH39* overexpression disappears in the absence of FIT, implying that bHLH Ib functions in a FIT dependent fashion. All four bHLH Ib members are significantly upregulated in the *fit* mutant (Wang et al., 2007), further supporting that FIT is required for bHLH Ib activating *IRT1* and *FRO2*. It is unclear whether FIT can function independently of bHLH Ib. Therefore, it is particularly of importance to clarify the molecular mechanism by which FIT and bHLH Ib TFs coordinate the expression of Fe uptake genes with the fluctuation of Fe availability.

Wang et al. (2013) revealed that the induction of *IRT1* and *FRO2* by Fe deficiency is decreased in the *bhlhl38 bhlh100 bhlh101* and *bhlhl39 bhlh100 bhlh101* mutants. In contrast, Sivitz et al. (2012) and Maurer et al. (2014) revealed that the expression of *IRT1* and *FRO2* is not affected in the *bhlh100 bhlh101* and *bhlhl39 bhlh100 bhlh101* mutants. Unlike *FIT* that is a root-specific gene, all four bHLH Ib genes are ubiquitously expressed both in the root and shoot under Fe deficiency conditions (Wang et al., 2007). A previous study concluded that bHLH Ib genes are involved in the leaf cell differentiation and chloroplast development (Andriankaja et al., 2014). It remains unclear whether bHLH Ib TFs have specific roles in regulating Fe deficiency response of shoots. Due to the functional redundancy of bHLH Ib members, functional clarification of bHLH Ib genes calls for the generation of their quadruple mutants. It is also an open question whether FIT and bHLH Ib have the identical contribution to the expression of Fe uptake genes.

In this study, we aimed to clarify whether FIT and bHLH Ib have the same functions in regulating Fe uptake and how they coordinate the expression of Fe uptake genes in the shoots and roots. We found that *fit* and *bhlh4x* (*bhlh38 bhlh39 bhlh100 bhlh101*) mutants have the identical Fe deficiency symptoms. Further transcriptome analysis revealed that they affect the expression of Fe uptake genes in the same way. Genetic evidence suggested that FIT and bHLH Ib function in the same genetic node. Further biochemical analysis found that bHLH Ib TFs without transcription activation activity have DNA binding activity and FIT without DNA binding activity has transcription activation activity. This work reveals that FIT and bHLH Ib form a functional transcription complex to activate the expression of Fe uptake genes.

## Results

### bHLH Ib quadruple mutants phenocopy the *fit* loss-of-function mutant

To avoid the functional redundancy between the four bHLH Ib members, we constructed the *bhlh38 bhlh39 bhlh100 bhlh101* (*bhlh4x*) quadruple mutants by editing *bHLH39* with CRISPR/Cas9 in the *bhlh38 bhlh100 bhlh101* background. When grown in soil, both *bhlh4x* and *fit-2* died at the seedling stage and this phenomenon could be rescued by extra Fe application (Figure 1A). The Fe concentration in *bhlh4x* was similar to that in *fit-2*, but significantly lower than that in wild type (Figure 1B). When grown on Fe deficient agar medium, *bhlh4x* and *fit-2* produced similar chlorotic leaves and short roots (Supplemental Figure S1A). In agreement with the chlorotic leaves, the chlorophyll concentration in both *bhlh4x* and *fit-2* was remarkably reduced (Supplemental Figure S1B). Next, we analyzed the H^+^-ATPase activity and ferric-chelate reductase activity, the typical indicators of Fe deficiency (Yi and Guerinot, 1996; Fox and Guerinot, 1998). In contrast to the visible coloration around the wild type roots, the coloration was hardly observed around the *bhlh4x* and *fit-2* roots (Figure 1C). Similarly, the ferric-chelate reductase activity was not increased significantly in both *bhlh4x* and *fit-2* under Fe deficient conditions (Figure 1D). Subsequently, we determined the expression of *IRT1* and *FRO2*, finding that their expression levels were similar between *bhlh4x* and *fit-2* (Figure 1E and F). Collectively, no matter under Fe sufficient or deficient conditions, the *bhlh4x* mutants completely mimicked *fit-2*.

**Figure 1.**
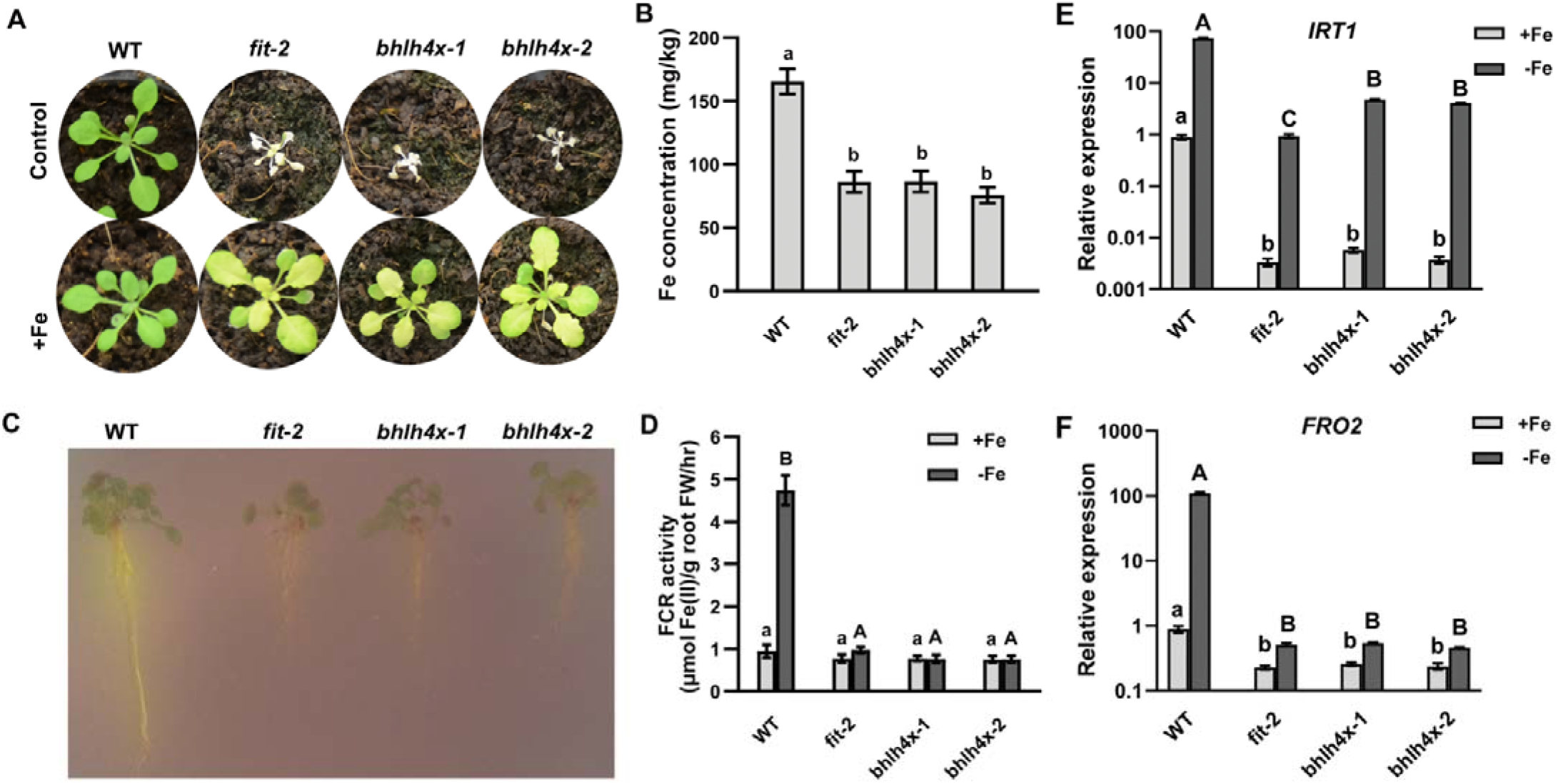
The Fe deficiency response of *fit-2* and *bhlh4x* mutants. (A) Phenotypes of *fit-2* and *bhlh4x*. Four-week-old plants are shown. ‘Control’ indicates that plants were watered with tap water; ‘+Fe’ indicates that plants were watered every three days with 0.5 mM Fe (II)-EDTA solution. (B) Fe concentration. Plants were watered every three days with 0.5 mM Fe (II)-EDTA solution. Leaves from four-week-old plants were used for Fe measurement. Data represent means ± standard deviation (SD) (*n* = 3). Different letters above each bar indicate statistically significant differences as determined by one-way ANOVA followed by Tukey’s multiple comparison test (P < 0.05). (C) Rhizosphere acidification. Seedlings grown on +Fe medium for 5 days were shifted to –Fe medium for 3 days, and then shifted to plates containing bromocresol purple. (D) FCR activity. Data represent means ± standard deviation (SD) (*n* = 3). Different letters above each bar indicate statistically significant differences as determined by one-way ANOVA followed by Tukey’s multiple comparison test. (E) and (F) Expression of *IRT1* (E) and *FRO2* (F) in *fit-2* and *bhlh4x*. Plants were grown on +Fe medium for 4 d and then transferred to +Fe or –Fe medium for 3 d. RNA was prepared from root tissues. Data represent means ± standard deviation (SD) (*n* = 3). The different letters above each bar indicate statistically significant differences as determined by one-way ANOVA followed by Tukey’s multiple comparison test (P < 0.05).

### FIT and bHLH Ib act in the same genetic node of Fe signaling pathway

To analyze how FIT and bHLH Ib TFs regulate the Fe deficiency response, we investigated transcriptomic changes in the *fit-2* and *bhlh4x*. Seven-day-old seedlings grown on Fe sufficient medium were shifted to Fe sufficient or deficient medium for 3 days, respectively. Shoots and roots were harvested separately for RNA sequencing. We identified 864 genes upregulated by Fe deficiency in a FIT dependent manner in roots, 66% of which also depended on bHLH Ib TFs (Figure 2A). Similarly, we identified 550 genes downregulated by Fe deficiency in a FIT dependent manner in roots, 60% of which also depended on bHLH Ib TFs (Figure 2A). Then, we focused on the expression of the well-known Fe deficiency responsive genes in roots. We found that the FIT dependent Fe-uptake associated genes, such as *IRT1, IRT2, FRO2, NICOTIANAMINE SYNTHASE 1* (*NAS1*), *NAS2, IRON REGULATED 2* (*IREG2*), *ZRT-AND IRT-RELATED PROTEIN 8* (*ZIP8*), *ZIP9, MYB10, MYB72, SCOPOLETIN 8-HYDROXYLASE* (*S8H), CYTOCHROME P450, FAMILY 82, SUBFAMILY C, POLYPEPTIDE 4* (*CYP82C4*), etc, were down-regulated in *fit*-2 and *bhlh4x*, whereas the FIT independent Fe deficiency responsive genes, such as *IRON MAN 1-4* (*IMA1-4*), *IMA6, FRO3, OLIGOPEPTIDE TRANSPORTER 3* (*OPT3*), *BRUTUS* (*BTS*), *POPEYE* (*PYE*), etc, were up-regulated in *fit-2* and *bhlh4x* (Figure 2B; Supplemental Table S1). These data suggest that bHLH Ib TFs regulate the Fe deficiency response of roots in a manner similar to FIT. Under Fe deficient conditions, *FIT* is mainly expressed in the root (Jakoby et al., 2004), and bHLH Ib TFs are abundant in both the root and shoot (Wang et al., 2007). However, the shoots of *bhlh4x* and *fit-2* displayed the identical phenotypes (Figure 1A; Supplemental Figure S1), and the typical Fe-deficiency responsive genes showed the very similar change trends in the shoots of *bhlh4x* and *fit-2* in response to Fe deficiency (Supplemental Figure S2; Supplemental Table S2). Given that FIT and bHLH Ib TFs affect the expression of Fe deficiency responsive genes in a similar fashion, we proposed that they act in the same genetic node of Fe deficiency response signaling pathway. To further confirm this hypothesis, we generated *bhlh4x-2 fit-2* quintuple mutants by crossing *bhlh4x-2* with *fit-2*. No matter in normal soil or in soil supplied with extra Fe, there is no visible difference between *bhlh4x-2, fit-2*, and *bhlh4x-2 fit-2* (Supplemental Figure S3). These results suggest that bHLH Ib and FIT play the same roles in Fe homeostasis.

**Figure 2.**
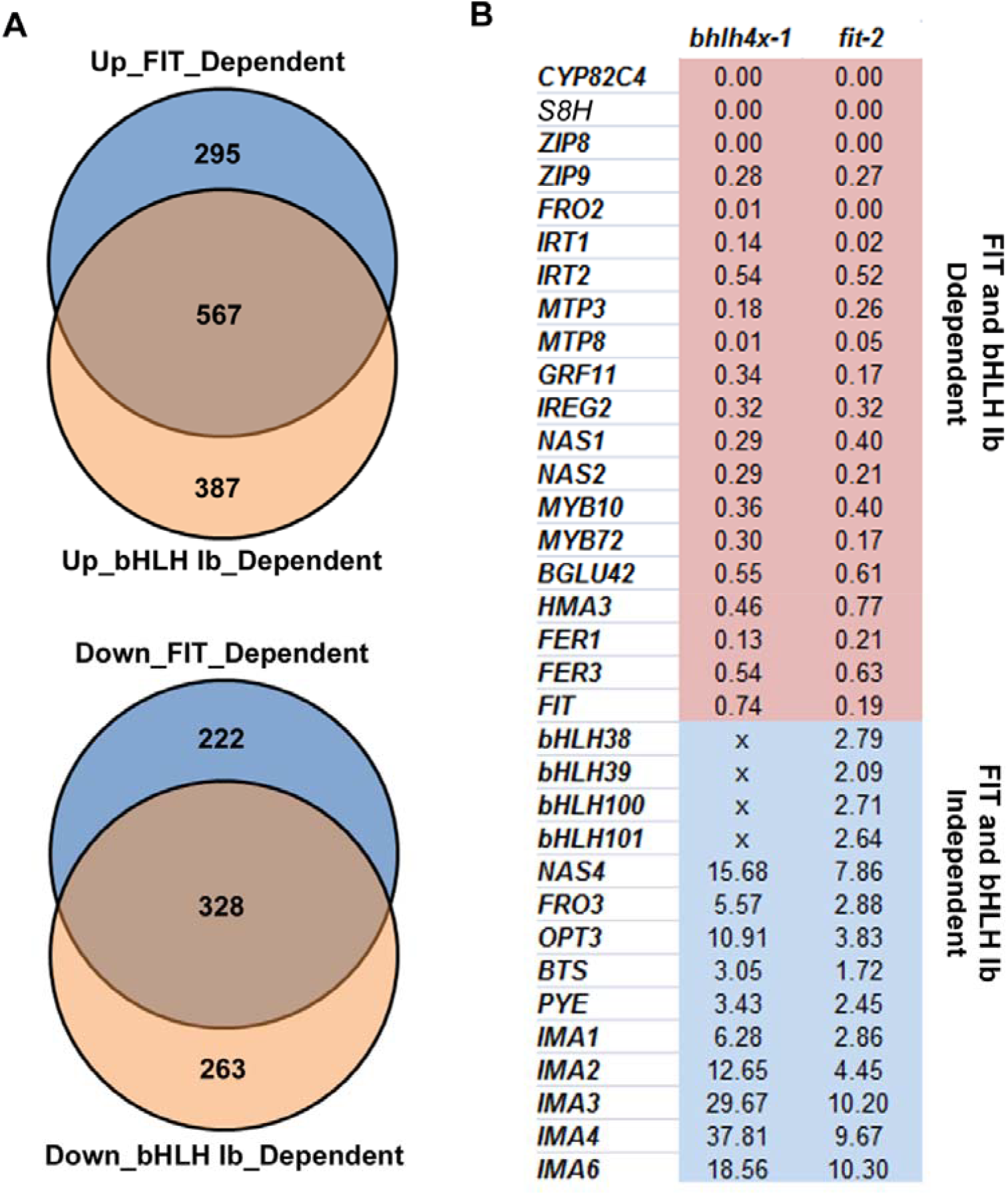
Expression of Fe deficiency responsive genes in the roots of *fit-2* and *bhlh4x-1*. (A) Venn diagram showing overlap between FIT-dependent and bHLH Ib dependent genes. (B)Relative transcript levels of genes involved in Fe deficiency response signaling in *fit-2* and *bhlh4x-1* under Fe-deficient conditions. The gene expression level in the wild type under Fe-deficient conditions was set to 1. Data are from the transcriptome in roots. “x” indicates that the RNA abundance for this gene is unavailable since the full-length CDS of *bHLH38/100/101* is undetectable and that of *bHLH39* is mutated.

### Overexpression of *FIT* cannot rescue *bhlh4x*

The four bHLH Ib members are significantly up-regulated in the *fit* mutant (Wang et al., 2007), implying that FIT is not required for the upregulation of bHLH Ib genes. The activation of *IRT1* and *FRO2* by *bHLH39* overexpression disappears in the absence of FIT (Naranjo-Arcos et al., 2017), implying that the functions of bHLH Ib TFs require the involvement of FIT. To further confirm whether the function of FIT requires bHLH Ib TFs, we generated the *FIT* overexpression plants (*bhlh4x*/*FIToe*) in the *bhlh4x-1* background. Irrespective of Fe status, the overexpression of *FIT* had no effect on the phenotypes of *bhlh4x-1*, as well as on the expression of *IRT1* and *FRO2* in the *bhlh4x-1* (Figure 3). Together, all these data suggest the functional interdependence between FIT and bHLH Ib in the Fe deficiency response.

**Figure 3.**
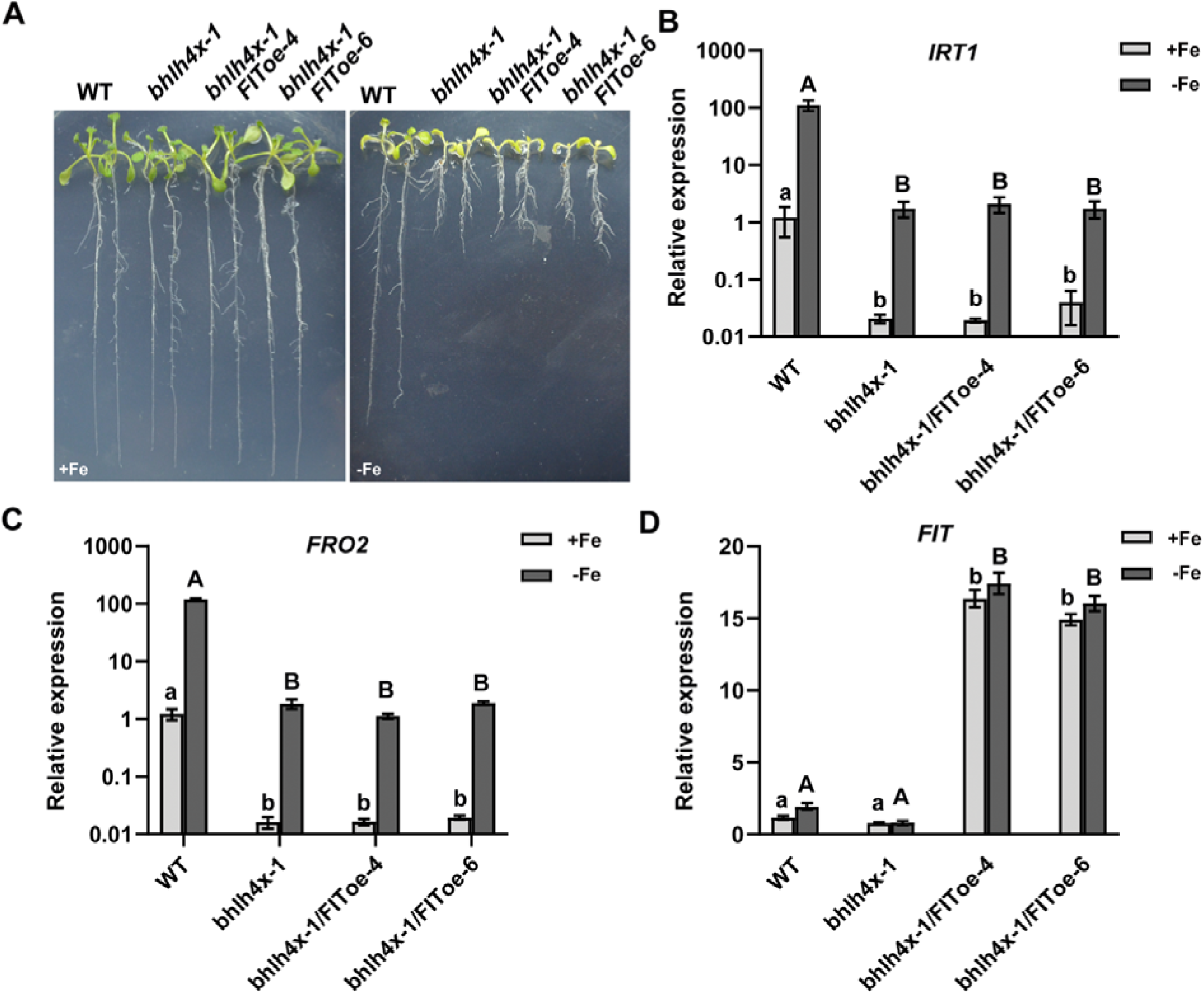
*FIT* overexpression does not rescue *bhlh4x-1*. (A) Phenotypes of 10-day-old seedlings grown on +Fe or –Fe medium are shown. (B-D) Expression of *IRT1* (B), *FRO2* (C) and *FIT* (D). Plants were grown on +Fe medium for 4 d and then transferred to +Fe or –Fe medium for 3 d. RNA was prepared from root tissues. Data represent means ± standard deviation (SD) (*n* = 3). The different letters above each bar indicate statistically significant differences as determined by one-way ANOVA followed by Tukey’s multiple comparison test (P < 0.05).

### Subcellular localization of bHLH Ib members

The phenotypic and genetic data support that FIT and bHLH Ib TFs depend on each other to function. Next, we explored why FIT and bHLH Ib require each other to function. A recent study revealed that bHLH39 moves to nuclei in a FIT dependent manner (Trofimov et al., 2019). Thus, it is likely that the bHLH Ib TFs need FIT to help them accumulate in nuclei and then activate Fe-uptake genes. To test this hypothesis, we fused the mCherry reporter to the C-end of bHLH Ib members and conducted transient expression assays (Figure 4A). Like bHLH39-mCherry, bHLH38-mCherry was mainly expressed in the cytoplasm. To determine the localization of bHLH38-mCherry in the presence of FIT, the GFP reporter was fused to the C-end of FIT. Unlike the GFP alone which was highly expressed in both the cytoplasm and nucleus, FIT-GFP was mainly expressed in the nucleus. The localization of bHLH38-mCherry did not change when co-expressed with the GFP. In contrast, most of bHLH38-mCherry was observed in the nucleus when co-expressed with the FIT-GFP (Figure 4B). This scenario is very similar to the case of bHLH39 (Trofimov et al., 2019). However, bHLH100-mCherry and bHLH101-mCherry were observed mainly in the nucleus in the absence of FIT (Figure 4A). Given the functional redundancy of bHLH Ib TFs and nuclear localization of bHLH100 and bHLH101 without the FIT assistance, we speculated that the FIT dependent nuclear accumulation of bHLH38 and bHLH39 is not the reason why bHLH Ib functionally depends on FIT.

**Figure 4.**
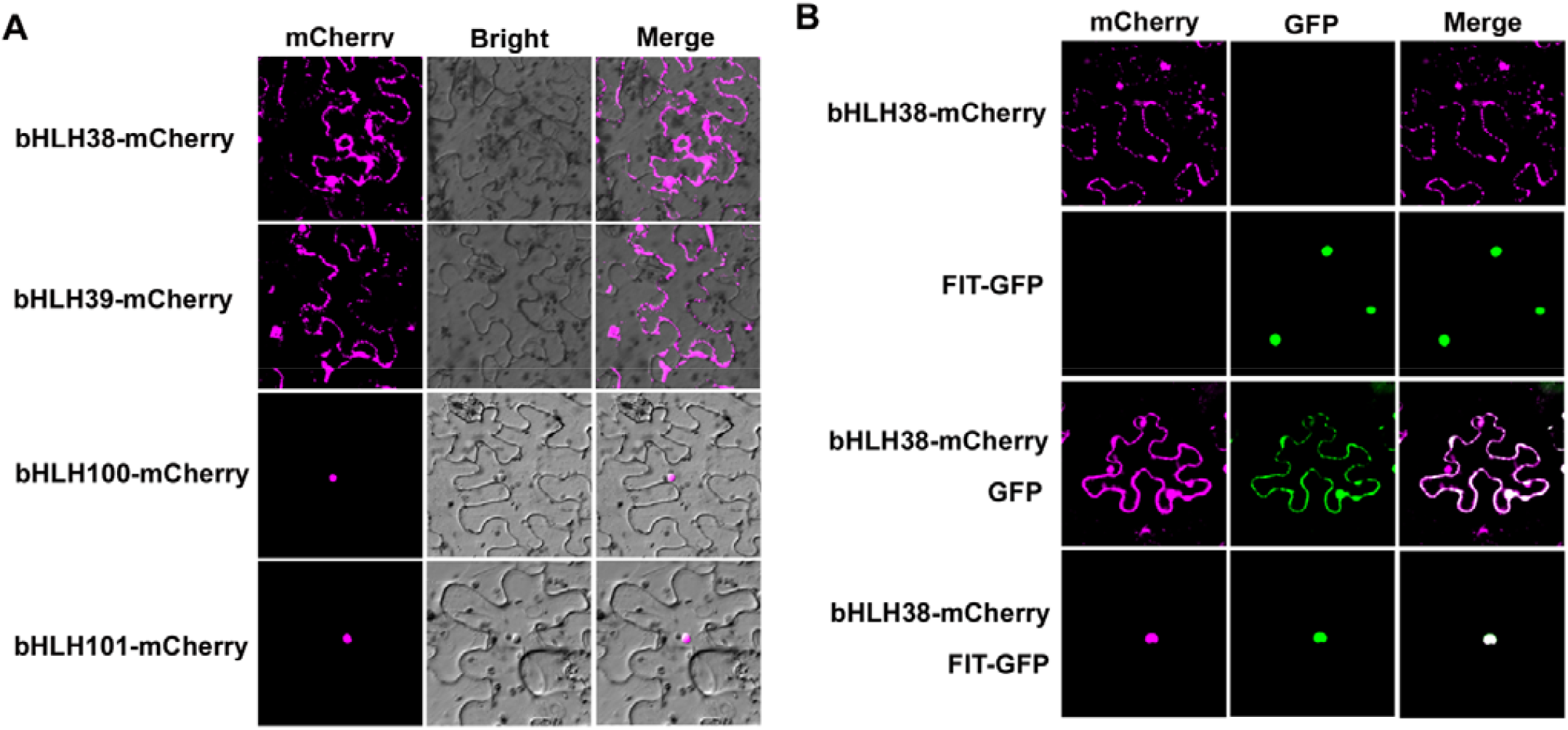
Subcellular localization of four bHLH Ib members. (A) Subcellular localization of bHLH Ib. bHLH38-mCherry, bHLH39-mCherry, bHLH100-mCherry, or bHLH101-mCherry, were expressed transiently in tobacco cells. (B)Effect of FIT on localization of bHLH38. bHLH38-mCherry, FIT-GFP, the combination of bHLH38-mCherry/GFP, and the combination of bHLH38-mCherry/FIT-GFP were expressed respectively in tobacco cells.

### bHLH Ib TFs have DNA binding ability, but no transactivation ability

Generally, eukaryotic TFs contain at least two domains, a DNA binding domain and a transcriptional activation or repression domain, which operate together to control the transcriptional initiation from target gene promoters. Subsequently, we wanted to know if FIT and bHLH Ib have the two characteristic domains. We employed an artificial GAL4 reporter system (Figure 5A), in which the yeast *GAL4* promoter (*pGAL4*) was used to drive a nuclear localization signal fused GFP (nGFP) as the reporter (*pGAL4:nGFP*) and the GAL4 DNA binding domain (BD) with a nuclear localization signal fused mCherry (nmCherry) was linked with a test protein X as the effector (BD-nmCherry-X). A strong transactivation domain VP16 from the herpes simplex virus was used as a positive control. Similar to the positive control, FIT activated the expression of *nGFP* whereas all four bHLH Ib TFs did not (Figure 5B). These data suggest that FIT, but not the four bHLH Ib TFs, has the transcriptional activation ability.

**Figure 5.**
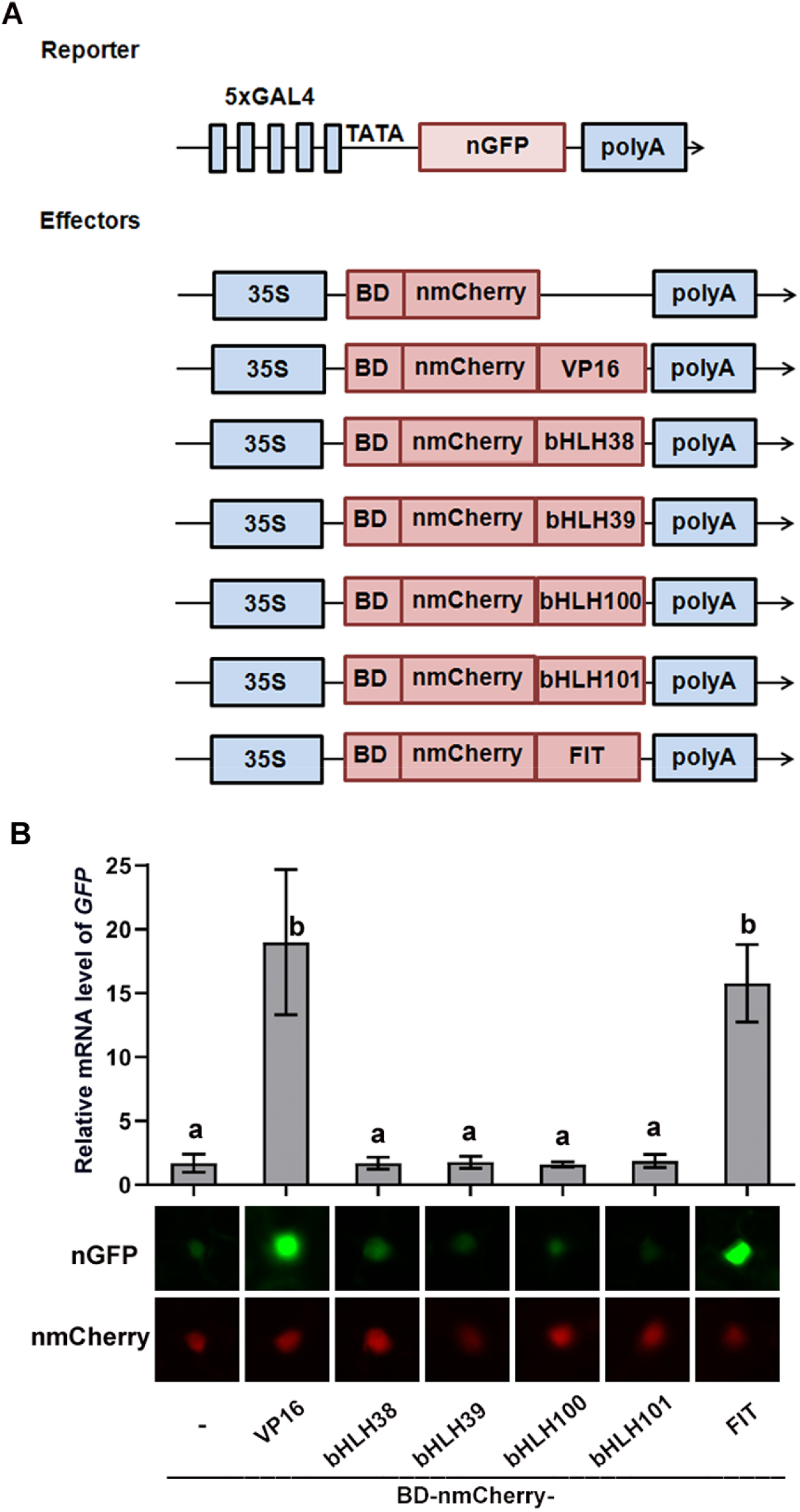
FIT has transcription activation ability. (A) Schematic diagram of constructs used in transient expression assays in(B) The reporter construct consists of five GAL4 binding motifs, a nuclear localization sequence tagged GFP (nGFP), and a poly(A) terminator. An effector consists of cauliflower mosaic virus 35S promoter (35S), GAL4 DNA binding domain (BD), a nuclear localization sequence tagged mCherry (nmCherry), a test gene, and a polyA terminator. VP16, an established transactivaion domain, was used as a positive control. (B) Determination of transactivation activity. Representative GFP and mCherry signals for each combination are shown. Expression of *nGFP* was normalized to *NPT II*. Data represent means ± standard deviation (SD) (*n* = 3). Different letters above each bar indicate statistically significant differences as determined by one-way ANOVA followed by Tukey’s multiple comparison test.

### FIT has transactivation ability, but no DNA binding ability

Subsequently, we wanted to know whether they have the DNA binding ability. It is well known that the bHLH domain of bHLH TFs is responsible for DNA binding and the amino acids at positions 5, 9, and 13 in the basic region are the most critical (Heim et al., 2003). The bHLH proteins with DNA binding ability have the conserved H/K-E-R residues at positions 5, 9, and 13 (Heim et al., 2003; Liu et al., 2013; Zhang et al., 2017, 2020; Lei et al., 2020). We aligned the bHLH domain of FIT and bHLH Ib, and found that the four Bhlh Ib TFs share an H-E-R motif, but FIT has a T-E-R motif (Figure 6A), implying that FIT may have no DNA binding ability. To provide experimental evidence, electrophoretic mobility shift assays (EMSA) were conducted. His-tagged bHLH38/39 and FIT were respectively expressed and purified from *Escherichia coli* and a fragment of *IRT1* promoter with an E-box was used the probe. The results suggest that bHLH38/39 can bind to the promoter of *IRT1* whereas FIT cannot (Figure 6B). Taken together, our data suggest that FIT has transcriptional activation ability and bHLH Ib has DNA binding ability. We propose that bHLH Ib and FIT form a functional transcription complex, in which bHLH Ib is responsible for target DNA binding and FIT for transcription activation.

**Figure 6.**
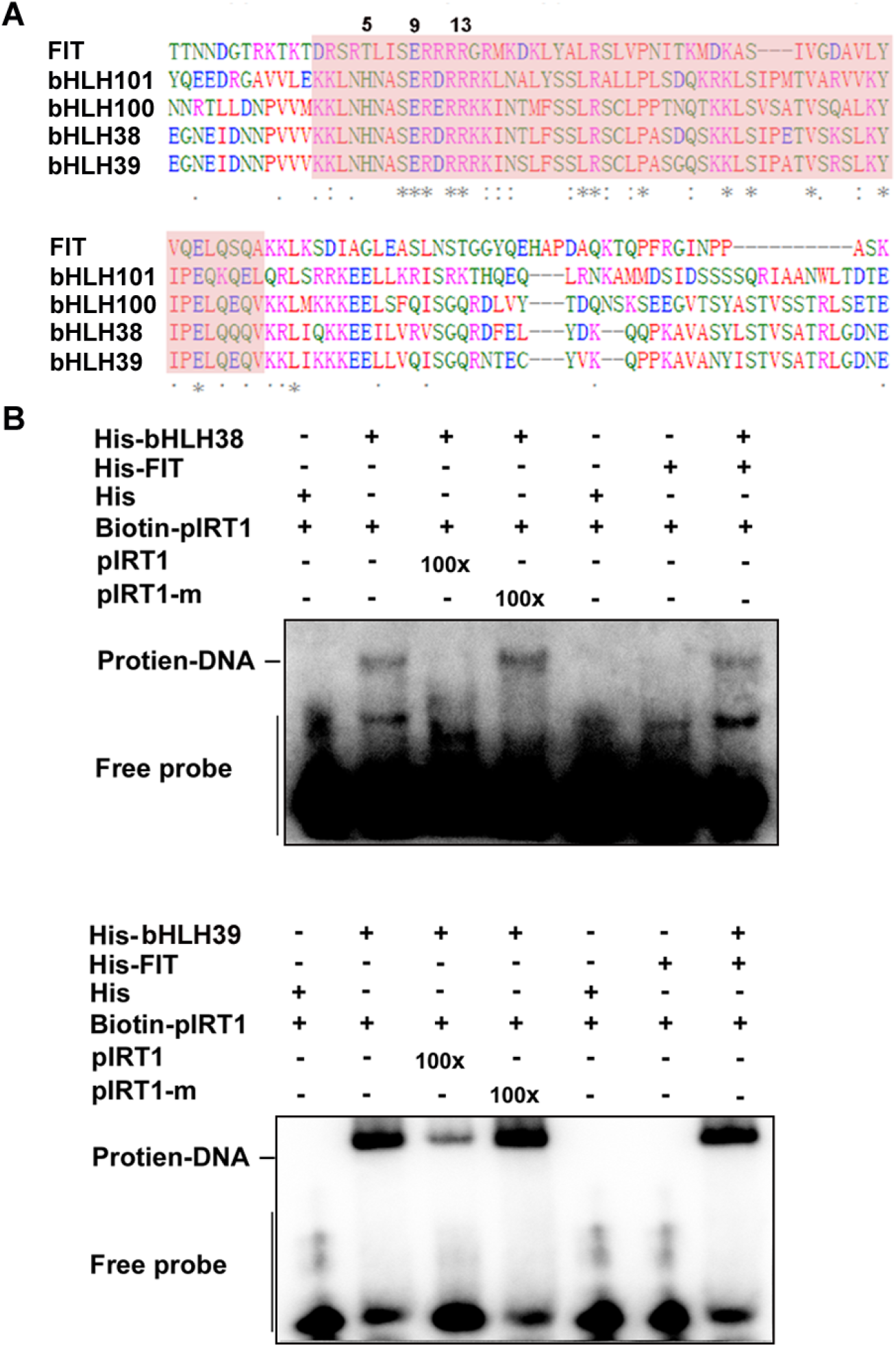
bHLH Ib TFs have DNA binding ability. (A)Alignment of bHLH domains of FIT and bHLH Ib. The regions in shadow represent the bHLH domains. The numbers (5, 9 and 13) indicate the 5^th^, 9^th^ and 14^th^ residues of bHLH domain. (B)EMSA assays. EMSAs were performed with a fragment of *IRT1* promoter. His-bHLH38, His-bHLH39, and His-FIT were used for binding assays. Biotin-probe, biotin-labeled probe; cold-probe, unlabeled probe; cold-probe-m, unlabeled mutated probe with mutated E-box.

### FIT and bHLH Ib form a functional transcription complex

We proposed that bHLH Ib and FIT form a functional transcription complex, in which bHLH Ib is responsible for target DNA binding and FIT for transcription activation. It has been confirmed that the activation of *IRT1* and *FRO2* by bHLH39 requires the involvement of FIT (Naranjo-Arcos et al., 2017). Given that FIT interacts with bHLH Ib and the latter directly binds to the promoter of *IRT1*, we wondered whether the FIT-bHLH Ib complex associates with the promoter of *IRT1 in vivo*. ChIP assays were performed in parallel among wild type, *FIToe* and *bhh4x-1/FIToe*. We observed FIT-specific enrichment for the *IRT1* promoter in the *FIToe* plants. However, this enrichment was dramatically reduced in the *bhh4x-1/FIToe* plants (Figure 7A). These data suggest that bHLH Ib is required for FIT association with target promoters.

**Figure 7.**
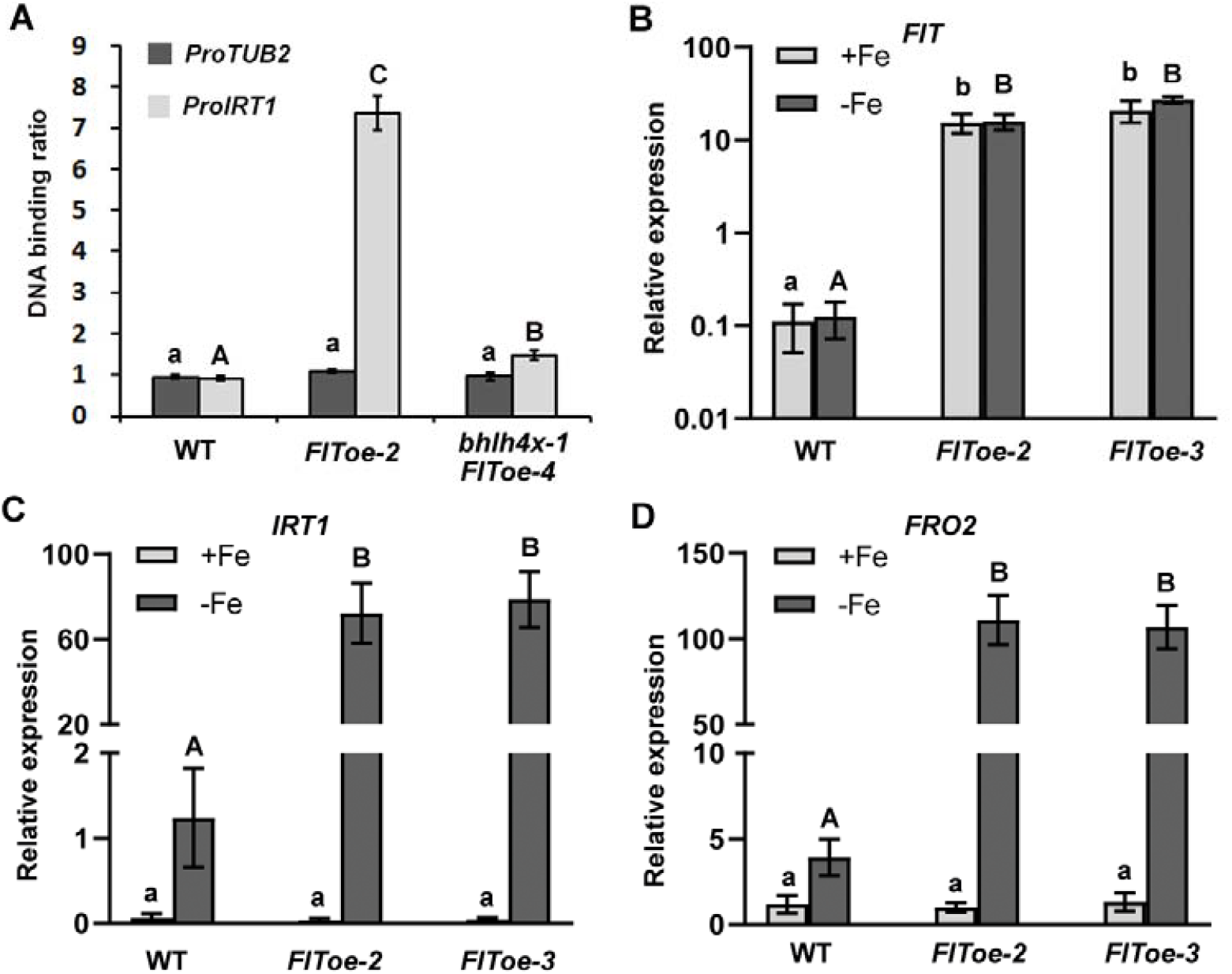
Association of FIT with the *IRT1* promoter requires bHLH Ib. (A)ChIP assays. Seven-day-old seedlings grown on +Fe media were shifted to -Fe media for three days. Whole seedlings were harvested for ChIP assays using anti-HA antibody, and the immunoprecipitated DNA was quantified by qPCR. The binding of the *TUB2* promoter fragment in the wild type was set to 1 and used to normalize the DNA binding ratio of the *IRT1* promoter. Data represent means ± SD (n = 3). The value which is significantly different from the corresponding control wild type was indicated by * (P<0.05), as determined by Student’ t test. (B)Expression of *IRT1* and *FRO2* in the leaves of *FIT* overexpression plants. Plants were grown on +Fe medium for 4 d and then transferred to +Fe or –Fe medium for 3 d. RNA was prepared separately shoots. Data represent means ± standard deviation (SD) (*n* = 3). The different letters above each bar indicate statistically significant differences as determined by one-way ANOVA followed by Tukey’s multiple comparison test (P < 0.05).

Although bHLH Ib genes are highly expressed in the shoots under Fe deficient conditions, the Fe-uptake genes (e. g. *IRT1*) are barely expressed. Having confirmed that FIT and bHLH Ib depend each other to initiate the transcription of their target genes, we wondered whether the expression of Fe-uptake genes would increase dramatically under Fe deficiency conditions if *FIT* was ectopically overexpressed in the shoots. To test this hypothesis, we determined the expression of *IRT1* and *FRO2* in the *FIT*oe plants (Figure 7B). In consistence with the high levels of *FIT* mRNA, the abundance of *IRT1* and *FRO2* was also at a considerably high level in the shoot of *FIT*oe under Fe deficient conditions. Collectively, these results support that FIT and bHLH Ib form a functional transcription complex to activate the expression of Fe uptake genes.

## Discussion

The transcription activation of Fe uptake genes and then Fe uptake is crucial for plants’ survival upon Fe deficiency conditions. FIT is a regulatory hub for iron deficiency and stress signaling in roots, which determines the expression of Fe uptake genes with the fluctuation of Fe status (Schwarz & Bauer, 2020). Two important questions regarding FIT and bHLH Ib are why the depend on each other to regulate Fe uptake, and whether they have their own independent functions in Fe signaling. Here, we address these two questions by physiological, genetic, and molecular evidence.

Although bHLH Ib genes are required for the expression of Fe uptake genes, their functional redundancy and lack of mutants with the loss-of-function of all four members make it hard to elucidate their exact contribution to Fe uptake. In terms of the Fe deficiency response, *fit-2* and *bhlh4x* mutants displayed the identical phenotypes as well as the same expression trends of Fe signaling associated genes, suggestive of the equal contribution of FIT and bHLH Ib to the Fe deficiency response. It was reported that the bHLH Ib member bHLH39 has no influence on the Fe-uptake genes in the absence of FIT (Naranjo-Arcos et al., 2017). We further confirmed that FIT cannot affect the Fe-uptake genes without bHLH Ib (Figure 3). These data suggest that FIT and bHLH Ib interdependently regulate the Fe-uptake genes.

Generally, a TF consists of a DNA binding domain and a transcription activation/repression domain. However, some TFs lost one of these two domains. For instance, IBH1 (ILI1 binding bHLH1) TF has no DNA binding ability, but interacts with ACEs (bHLH transcriptional activators for cell elongation) and interferes with the DNA binding ability and the transcription activation activity of the latter (Ikeda et al., 2012). In Arabidopsis, several single-repeat R3-MYB TFs, such as TRY (Triptychon), CPC (Caprice), ETCs (Enhancer of TRY and CPC 1, 2, 3), act as negative regulators of trichome differentiation, which contain a single DNA binding domain, but lack the activation domain (Wang and Chen, 2014). Here, we show that FIT has no DNA binding domain and bHLH Ib no transcription activation domain, and they both complement each other to form a functional transcription complex to initiate the transcription of their target genes. This two-component model (Figure 8) rationally explains why the overexpression of one of both cannot activate their targets in the absence of the other (Figure S4; Naranjo-Arcos et al., 2017) and why *bhlh4x* phenocopies *fit* (Figure 1; Figure S1).

**Figure 8.**
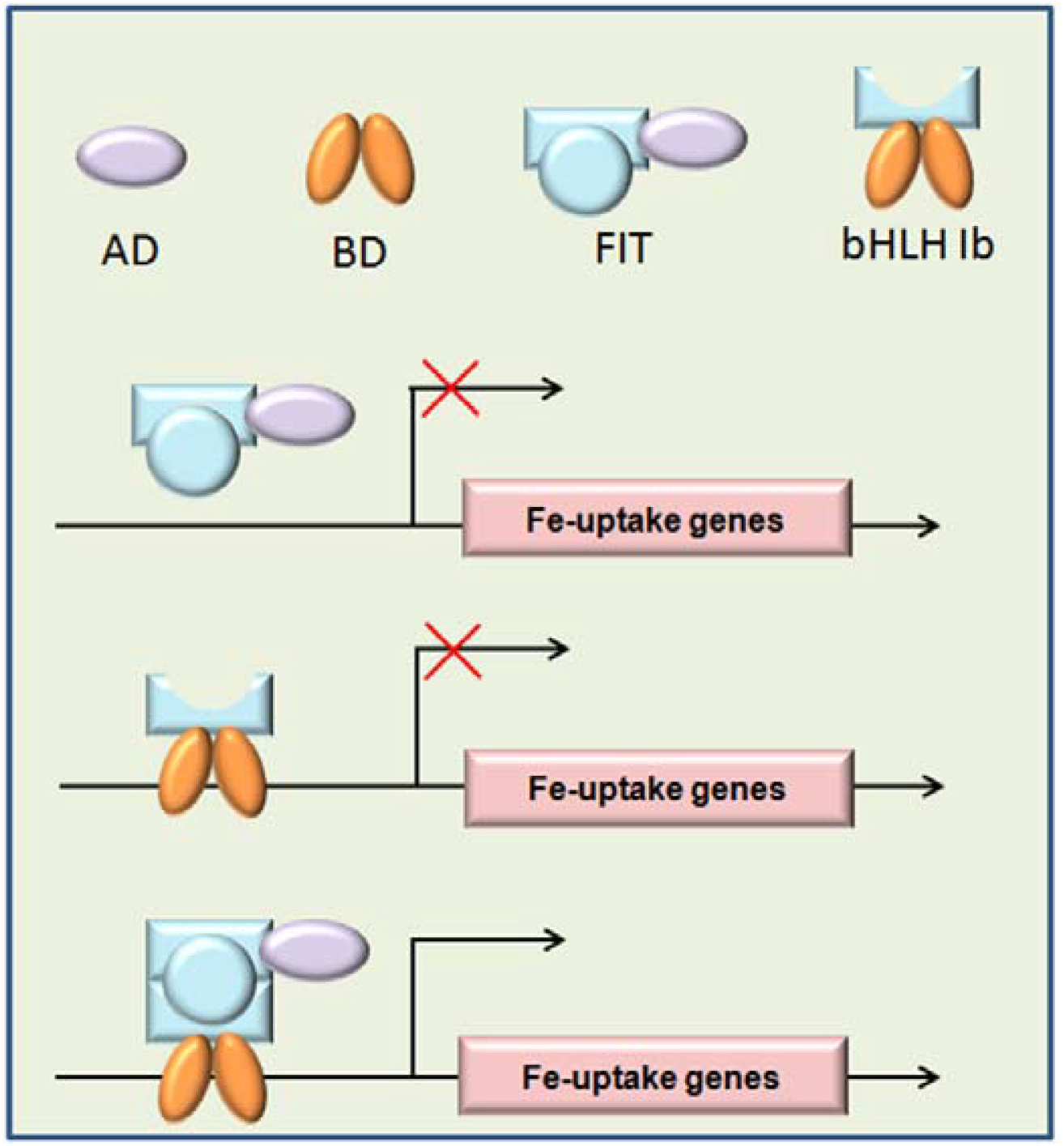
A proposed working model of FIT and bHLH Ib. bHLH Ib has a DNA binding domain (BD) responsible for target recognition. FIT has a transcription activation domain (AD) responsible for transactivation. FIT and bHLH Ib form a functional transcriptional complex. FIT alone cannot bind to the promoters of Fe uptake genes, and bHLH Ib alone cannot initiate the transcription of Fe-uptake genes. The combination of FIT and bHLH Ib results in a functional complex to activate the transcription of Fe-uptake genes.

It is well known that the expression of Fe-uptake genes changes spatio-temporally in response to Fe status. As the positive regulators of Fe-uptake genes, bHLH Ib TFs are ubiquitously expressed in the roots and shoots in response to Fe deficiency. In contrast, the Fe-uptake genes (e. g. *IRT1* and *FRO2*) are highly expressed in the Fe deficient roots, but hardly expressed in the Fe deficient shoots. The working model of FIT and bHLH Ib gives a reasonable explanation to the differential expression patterns of Fe uptake genes and bHLH Ib. In the Fe deficient shoots, although the transcript abundance of bHLH Ib is at a high level, that of *FIT* is at a low level, hence resulting in less FIT-bHLH Ib dimmers and then less Fe-uptake genes. Under Fe sufficient conditions, the Fe-uptake genes are not activated in the roots because bHLH Ib genes are expressed at a low level. Therefore, FIT is the limitation factor for Fe-uptake genes in the shoot, and bHLH Ib is the limitation factor in the roots.

The previous studies have revealed that bHLH Ib TFs regulate cell differentiation and chloroplast development (Andriankaja et al., 2014), in agreement with their expression in leaves. In contrast, *FIT* is barely expressed in leaves irrespective of Fe status. We noted that both *fit-2* and *bhlh4x* mutants produced leaves with serrate margin (Figure 1A), implying that FIT might also function in leaf development. Four bHLH Ib genes are involved in Fe uptake in Arabidopsis whereas only one (OsIRO2, IRON-RELATED BHLH TRANSCRIPTION FACTOR 2) in rice. bHLH38/39 are mainly localized in the cytoplasm and bHLH100/101 in the nucleus. Similar to bHLH100/101, OsIRO2 is also preferentially localized in the cytoplasm. Interestingly, the cytoplasm localized bHLH Ib proteins accumulate in the nucleus in the presence of their interaction partner FIT (Figure 3; Trofimov et al., 2019; Liang et al., 2020; Wang et al., 2020). Further investigation is needed to clarify whether the property of cytoplasmic localization is required for the molecular functions of bHLH Ib proteins.

In summary, the genetic and molecular data presented in this study suggest that the physical interaction of FIT and bHLH Ib leads to the formation of a functional transcription activation complex in which bHLH Ib exerts the DNA binding function and FIT exerts the transactivation function (Figure 8). This model explains why FIT and bHLH Ib depend on each other to activate the expression of Fe uptake genes, providing insights into the molecular mechanism by which plants control the expression of Fe-uptake genes in response to Fe deficiency.

## Materials and Methods

### Plant materials and growth conditions

*Arabidopsis thaliana* ecotype Col-0 was used as the wide-type. Seeds were surface-sterilized with 20% commercial bleach for 15 min and then washed three times with distilled water. After plated on half MS media, seeds were vernalized for 2 d at 4°C before germination in greenhouse. The ordinary medium was the half MS with 1% sucrose, 0.7% agar A, 0.1 mM Fe-EDTA at pH 5.8. For Fe deficiency media, the same half MS without Fe-EDTA was used. Plates were placed in a culture room at 22°C under a 16 h light/8 h dark photoperiod. *fit-2* (SALK_126020), *bhlh4x-1* and *bhlh4x-2* were described previously (Cai et al., 2021). The identification of *bhlh4x-1 fit-2* is shown in Supplemental Figure S3.

### High-throughput sequencing of mRNA, and differential gene expression analysis

For global analysis of gene expression, roots of wild-type, *bhlh4x-1* and *fit-2* plants grown on +Fe for 4 days and transferred to –Fe for 3 days. Roots and shoot were separately harvested and collected in liquid nitrogen. Total RNA of each sample was extracted according to the instruction manual of the TRlzol Reagent (Life technologies, California, USA). RNA integrity and concentration were checked using an Agilent 2100 Bioanalyzer (Agilent Technologies, Inc., Santa Clara, CA, USA). The mRNA was isolated by NEBNext Poly (A) mRNA Magnetic Isolation Module (NEB, E7490). The cDNA library was constructed following the manufacturer’s instructions of NEBNext Ultra RNA Library Prep Kit for Illumina (NEB, E7530) and NEBNext Multiplex Oligos for Illumina (NEB, E7500). In briefly, the enriched mRNA was fragmented into approximately 200nt RNA inserts, which were used to synthesize the first-strand cDNA and the second cDNA. The double-stranded cDNA was performed end-repair/dA-tail and adaptor ligation. The suitable fragments were isolated by Agencourt AMPure XP beads (Beckman Coulter, Inc.), and enriched by PCR amplification. Finally, the constructed cDNA libraries were sequenced on an Illumina sequencing platform. The raw sequencing data are stored in NCBI under the accession number of PRJNA694484 (https://www.ncbi.nlm.nih.gov/sra/PRJNA694484).

The values of fragments per kilobase of transcript per million mapped reads (FPKM) are shown for each gene. The genes for which no hits were recorded across all the samples were discarded from the data set. For the genes whose hits were recorded in only a subset of the samples, we replaced missing values with a small value of expression (0.01 FPKM). Transcript abundance was concluded to increase/decrease under Cu deficiency for a gene when arithmetic means of transcript abundance differed by a factor of at least 2. Changes in transcript levels were concluded to be dependent on FIT if log_2_ FC (wild type -Fe versus fit -Fe) > 1 for log_2_ FC (wild type -Fe versus wild type +Fe) > 1, and log_2_ FC (wild type -Fe versus fit -Fe) < 1 for log_2_ FC (wild type -Fe versus wild type +Fe) < -1. The bHLH Ib dependent transcripts were obtained by a similar filtration.

### Transient expression assays

All related plasmids were transformed into *Agrobacterium tumefaciens* strain EHA105. Agrobacterial cells were infiltrated into leaves of *Nicotiana benthamiana* by the infiltration buffer (0.2 mM acetosyringone, 10 mM MgCl_2_, and 10 mM MES, pH 5.6). In the transient expression assays, the final optical density at 600 nm value was 0.5 (reporter, pGAL4-nls-GFP) and 0.5 (effector, BD-nmCherry-X). After infiltration 2 days in dark, GFP fluorescence were observed through a Carl Zeiss Microscopy GmbH.

### EMSA

bHLH38, bHLH39 and FIT were respectively cloned into the pET-28a(+) vector and the resulting plasmids were introduced into *Escherichia coli* BL21(DE3) for protein expression. Cultures were incubated with 0.5 M isopropyl β-D-1-thiogalactopyranoside at 22°C for 16h, and proteins were extracted and purified by using the His-tag Protein Purification Kit (Beyotime, China) following the manufacturer’s protocol. EMSA was performed using the Chemiluminescent EMSA Kit (Beyotime, China). For generation of competitive probe (pIRT1) or mutated probe (pIRT1-m), a pair of complementary single-strand DNA primers were synthesized. For generation of the biotin-labeled probe (Biotin-pIRT1), a pair of complementary single-strand DNA primers with a biotin label at the 5’ end were synthesized. A pair of complementary primers were used for annealing to form double-strand DNA. The annealing reaction solution for 1 X probe was as follow: 1 μl of 10 μM forward prime, 1 μl of 10 μM reverse primer, 3 μl of 10 X Taq buffer, and 25 μl of H_2_O. The annealing reaction solution for 100 X probe was as follow: 10 μl of 100 μM forward prime, 10 μl of 100 μM reverse primer, 3 μl of 10 X Taq buffer, and 7 μl of H_2_O. Reaction solution was incubated at 95°C for 2 minutes, and cool at room temperature. The binding reaction solution was as follow: 5 μl of H_2_O, 2 μl of 5 X EMSA/Gel-Shift binding buffer, 2 μl of protein, 1 μl of probe. The competitive binding reaction solution was as follow: 4 μl of H_2_O, 2 μl of 5 X EMSA/Gel-Shift binding buffer, 2 μl of protein, 1 μl of 1 X probe, and 1 μl of 100 X probe. Incubate binding reactions at room temperature for 20 minutes. Add 1 μl of 10 X Loading Buffer to each 10 μl binding reaction, pipetting up and down several times to mix. Electrophorese binding reactions in a 6% polyacrylamide gel. Electrophoretic transfer in 0.5 X TBE at 380 mA (∼100 V) for 30 minutes. After crosslinking at 120 mJ/cm^2^, the membrane was incubated in 15 ml of Blocking Buffer for 15 minutes with gentle shaking. Then, the membrane was shifted to 15 ml of Blocking Buffer with 7.5 μl of Streptavidin-HRP Conjugate for 15 minutes with gentle shaking. Transfer membrane to a new container and rinse it briefly with 20 ml of 1 X wash solution. Wash membrane four times for 5 minutes each in 20 ml of 1 X wash solution with gentle shaking. Transfer membrane to 20 ml of Substrate Equilibration Buffer. Incubate membrane for 5 minutes with gentle shaking. Transfer membrane to 5 ml of Substrate Working Solution for 5 minutes. Expose membrane to a low-light cooled CCD imaging apparatus (Tanon-5200).

### Gene expression analysis

Total RNA was extracted using the RNAplant (Real-Times, China). cDNA was synthesized by the use of PrimeScript™RT reagent Kit with gDNA Eraser (Perfect Real Time) according to the reverse transcription protocol (Takara). The resulting cDNA was subjected to relative quantitative PCR using a SYBR Premix Ex Taq™ kit (TaKaRa) on a Roche LightCycler 480 real-time PCR machine, according to the manufacturer’s instructions. The relative expression of genes was normalized to that of *ACT2* and *PP2A*.

### Plasmid construction and generation of transgenic plants

For construction of transient expression vectors, the GAL4 binding domain was fused with mCherry containing a nuclear localization signal to generate 35S:BD-nmCherry. VP16, bHLH38, bHLH39, bHLH100 bHLH101 and FIT were respectively fused with BD-nmCherry as the effectors. The pGAL4 promoter driving a GFP containing a nuclear localization signal was as the reporter. For the subcellular localization assays, bHLH38, bHLH39, bHLH100 and bHLH101 were fused with mCherry respectively, and FIT was fused with GFP. For the construction of *bhlh4x-1/FIToe* plants, the HA-tagged FIT driven by 35S promoter was introduced into *bhlh4x-1* mutant by transformation.

### Chromatin immunoprecipitation (ChIP) assays

ChIP assays were conducted according to previously described protocols (Saleh et al., 2008). Plants grown on +Fe media for 7 days were shifted to –Fe media for 3 days, and then whole seedlings were used for ChIP assays. To quantify FIT-DNA binding ratio, qPCR was performed with the *TUB2* as the endogenous control.

### Subcellular localization

The full-length FIT was fused with GFP to generate FIT-GFP and the full-length bHLH38/39/100/101 with mCherry to generate bHLH38/39/100/101-mCherry. The plasmids above were transformed into agrobacteria. Agrobacteria were incubated in LB liquid media. When growth reached an OD600 of approximately 3.0, the bacteria were spun down gently (3200 g, 5 min), and the pellets were resuspended in infiltration buffer (10 mM MgCl_2_, 10 mM MES, pH 5.6) at a final OD600 of 1.0. A final concentration of mM acetosyringone was added. The agrobacteria were kept at room temperature for at least 2 h without shaking. Leaf infiltration was conducted in 3-week-old *N. benthamiana*. Excitation laser wave lengths of 488 nm and 563 nm were used for imaging GFP and mCherry signals, respectively.

## Acknowledgments

We thank the Biogeochemical Laboratory and Central Laboratory (Xishuangbanna Tropical Botanical Garden) for assistance in the determination of Fe concentration. We thank Germplasm Bank of Wild Species in Southwest China for confocal laser scanning microscopy.

## SConflict of interest statement

None declared.

## Supporting Information

Additional Supporting Information may be found online in the Supporting Information section at the end of the article.

**Supplemental Figure S1**. Comparison of *fit-2* and *bhlh4x* mutants.

**Supplemental Figure S2**. Transcripts responsive to Fe Deficiency in a FIT or bHLH Ib dependent fashion in shoots.

**Supplemental Figure S3**. Phenotypes of *bhlh4x-2 fit-2* quintuple mutants.

**Supplemental Figure S4**. Expression of *IRT1* and *FRO2* in the roots of *FIT* overexpression plants.

**Supplemental Table S1. Expression of Fe-deficiency responsive genes in roots**.

**Supplemental Table S2. Expression of Fe-deficiency responsive genes in shoots**.

**Supplemental Table S3**. Primers used in this paper.

