## Supplementary data for "Why FIT and bHLH Ib interdependently regulate Fe-uptake"

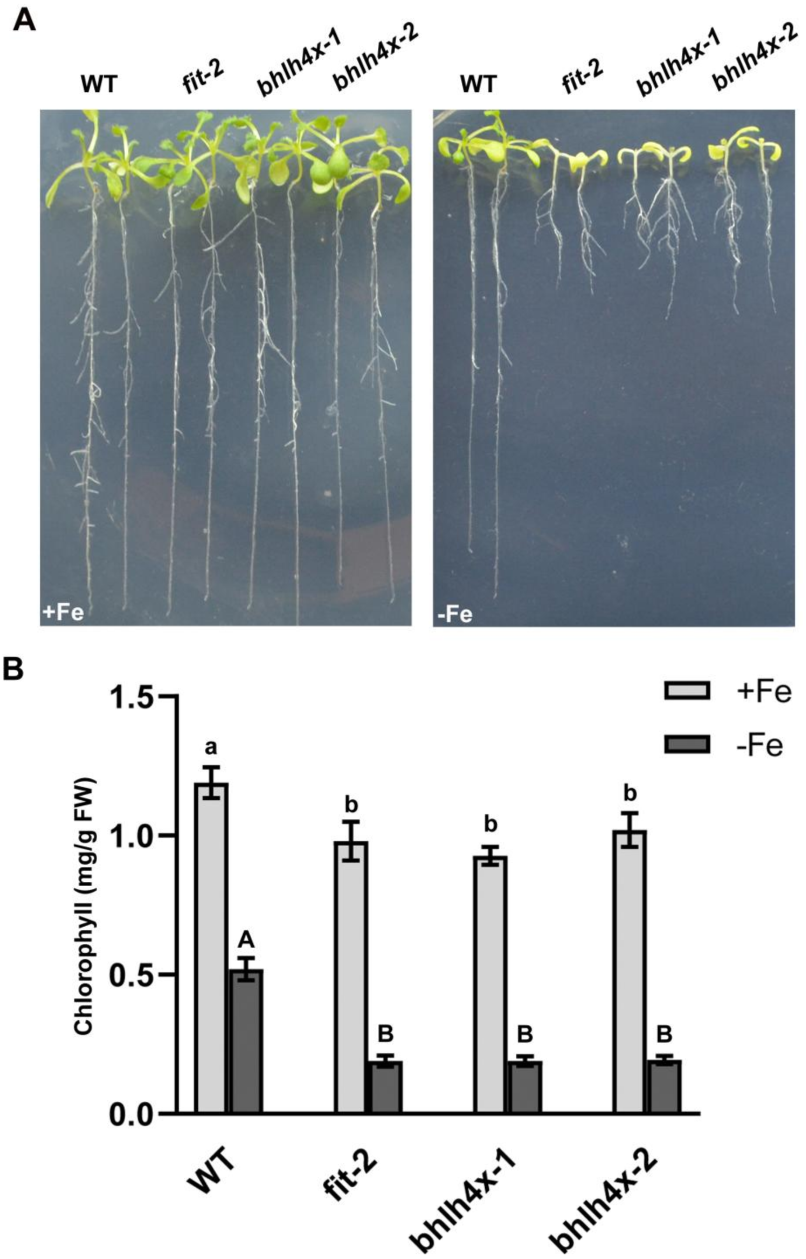


**Supplemental Figure S1**. Comparison of *fit-2* and *bhlh4x* mutants.

(A) Ten-day-old seedlings grown on +Fe or –Fe medium are shown.

(B) Seedlings grown on +Fe or –Fe medium for two weeks were used for chlorophyll measurement. The different letters above each bar indicate statistically significant differences as determined by one-way ANOVA followed by Tukey's multiple comparison test ($P < 0.05$).

A

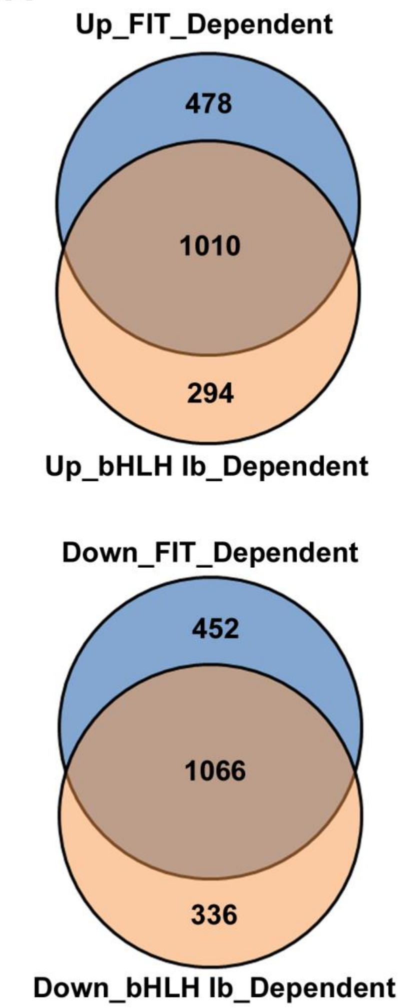


B

| | *bhlh4x-1* | *fit-2* | |
|---|---|---|---|
| ***CYP82C4*** | 0.02 | 0.02 | FIT and bHLH Ib dependent |
| ***S8H*** | 0.03 | 0.03 | |
| ***NAS2*** | 0.01 | 0.05 | |
| ***IREG2*** | 0.69 | 0.15 | |
| ***MTP3*** | 0.75 | 0.72 | |
| ***IRT1*** | 0.31 | 0.18 | |
| ***FER1*** | 0.58 | 0.56 | |
| ***FER3*** | 0.51 | 0.69 | |
| ***FIT*** | 0.16 | 0.30 | |
| ***bHLH38*** | x | 3.21 | FIT and bHLH Ib independent |
| ***bHLH39*** | x | 4.61 | |
| ***bHLH100*** | x | 4.42 | |
| ***bHLH101*** | x | 6.41 | |
| ***PYE*** | 2.01 | 1.39 | |
| ***BTS*** | 3.76 | 3.24 | |
| ***OPT3*** | 3.55 | 2.41 | |
| ***FRO2*** | 6.37 | 2.48 | |
| ***FRO3*** | 2.50 | 1.47 | |
| ***NAS4*** | 2.50 | 2.03 | |
| ***IMA1*** | 4.01 | 2.30 | |
| ***IMA2*** | 4.28 | 2.50 | |
| ***IMA3*** | 2.54 | 1.99 | |
| ***IMA4*** | 8.07 | 2.67 | |
| ***IMA6*** | 10.71 | 3.91 | |

**Supplemental Figure S2**. Transcripts responsive to Fe Deficiency in a FIT or bHLH Ib dependent fashion in shoots.

(A) Venn diagram showing overlap between FIT-dependent and bHLH Ib dependent genes.

(B) Expression of genes involved in Fe deficiency response signaling in *fit-2* and *bhlh4x-1* under Fe deficiency conditions. The gene expression level in the wild type under Fe-deficient conditions was set to 1. Data are from the transcriptome in shoots. “x” indicates that the RNA abundance for this gene is unavailable since the full-length CDS of *bHLH38/100/101* is undetectable and *bHLH39* is mutated (Supplemental Figure S3).

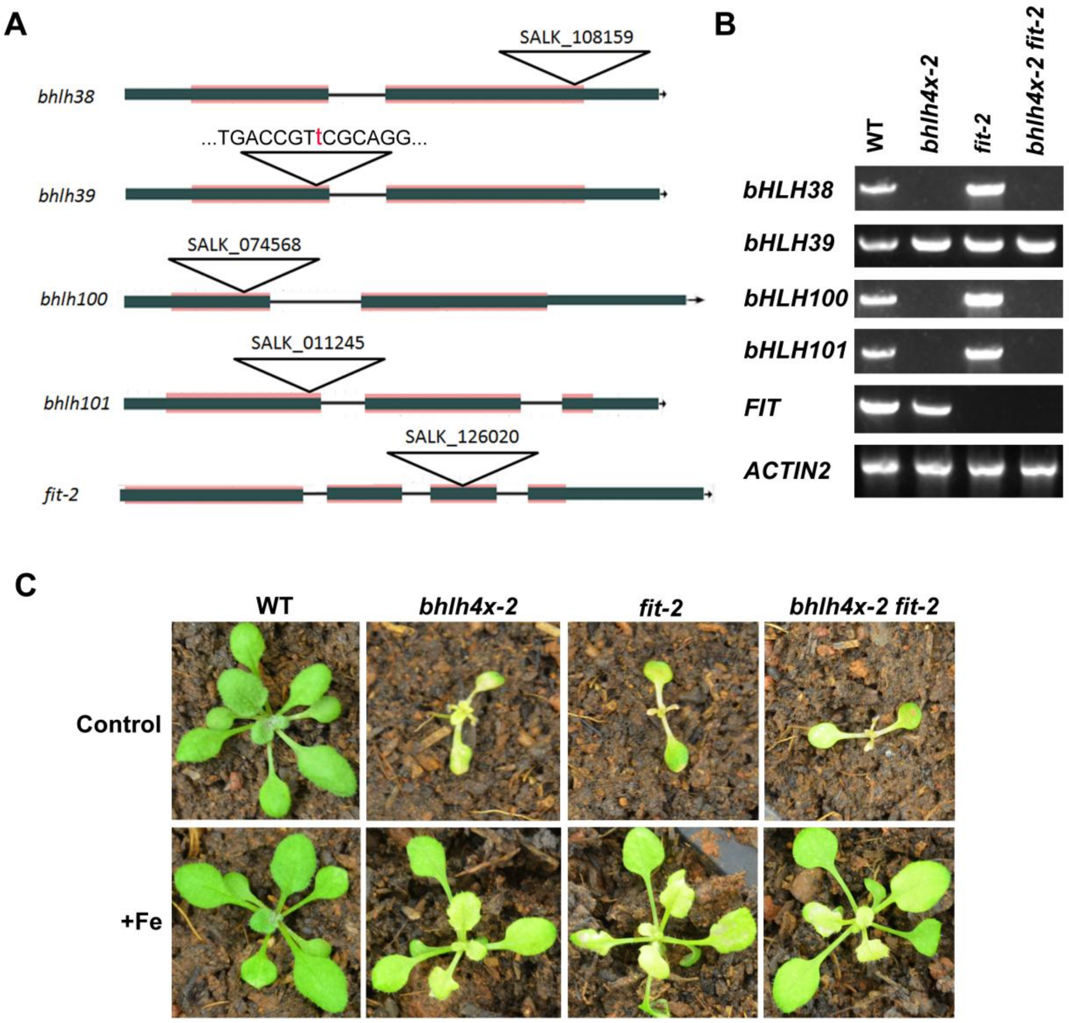


**Supplemental Figure S3**. Phenotypes of *bhlh4x-2 fit-2* quintuple mutants.

(A) The genotype of each mutated gene in the quintuple mutants. The red mask indicates the coding region. The red letter "t" indicates the inserted nucleotide in the *bHLH39* gene.

(B) Identification of *bhlh4x-2 fit-2* quintuple mutants. The full-length CDS for each gene was amplified by RT-PCR.

(C) Four-week-old plants are shown. 'Control' indicates that plants were watered with tap water; '+Fe' indicates that plants were watered every three days with 0.5 mM Fe(II)-EDTA solution.

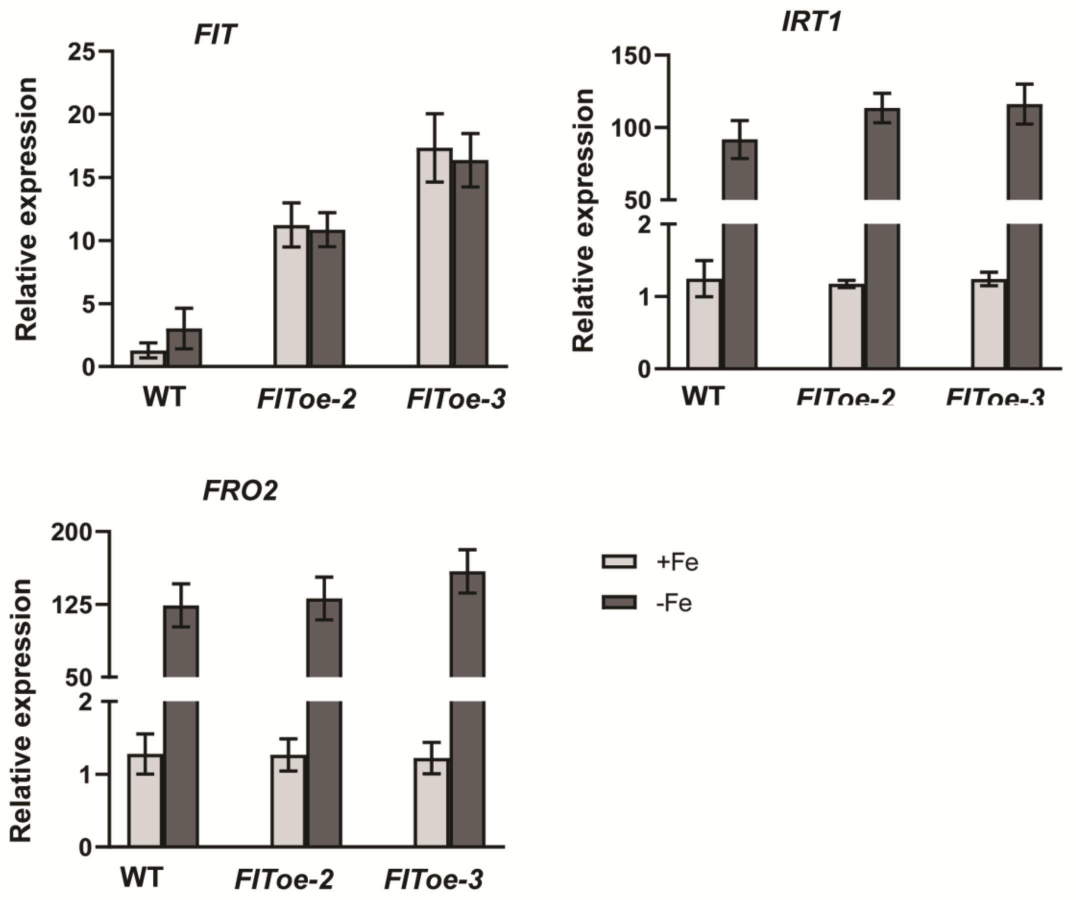


**Supplemental Figure S4**. Expression of *IRT1* and *FRO2* in the roots of *FIT* overexpression plants.

Plants were grown on +Fe medium for 4 d and then transferred to +Fe or –Fe medium for 3 d. RNA was prepared separately roots. Data represent means ± standard deviation (SD) (*n* = 3). The different letters above each bar indicate statistically significant differences as determined by one-way ANOVA followed by Tukey's multiple comparison test (P < 0.05).

**Supplemental Table S1.** Expression of Fe-deficiency responsive genes in roots.

| ID | AGI | +Fe | | | -Fe | | |
|---|---|---|---|---|---|---|---|
| | | WT | *bhlh4x-1* | *fit-2* | WT | *bhlh4x-1* | *fit-2* |
| *CYP82C4* | AT4G31940 | 0.55 | 0.01 | 0.01 | 125.55 | 0.01 | 0.01 |
| *S8H* | AT3G12900 | 0.05 | 0.05 | 0.01 | 71.18 | 0.07 | 0.01 |
| *ZIP8* | AT5G45105 | 0.01 | 0.01 | 0.01 | 3.26 | 0.01 | 0.01 |
| *ZIP9* | AT4G33020 | 0.3 | 1.15 | 0.43 | 1.97 | 0.55 | 0.53 |
| *FRO2* | AT1G01580 | 0.94 | 0.38 | 0.22 | 103.03 | 0.78 | 0.41 |
| *IRT1* | AT4G19690 | 5.63 | 3.26 | 0.09 | 213.58 | 30.54 | 3.38 |
| *IRT2* | AT4G19680 | 5.61 | 16.51 | 4.47 | 15.81 | 8.47 | 8.25 |
| *MTP3* | AT3G58810 | 6.2 | 9.19 | 7.5 | 30.98 | 5.57 | 8.2 |
| *MTP8* | AT3G58060 | 0.06 | 0.5 | 0.01 | 2.62 | 0.03 | 0.13 |
| *GRF11* | AT1G34760 | 1.71 | 3.95 | 1.2 | 7.73 | 2.63 | 1.3 |
| *IREG2* | AT5G03570 | 5.58 | 7.15 | 3.66 | 9.44 | 2.99 | 2.99 |
| *NAS1* | AT5G04950 | 48.15 | 120.84 | 71.45 | 138.93 | 40.7 | 55.03 |
| *NAS2* | AT5G56080 | 19.64 | 38.76 | 11.59 | 32.56 | 9.45 | 6.87 |
| *MYB10* | AT3G12820 | 3.65 | 2.22 | 2.47 | 8.57 | 3.05 | 3.41 |
| *MYB72* | AT1G56160 | 0.01 | 0.01 | 0.01 | 1.97 | 0.6 | 0.33 |
| *BGLU42* | AT5G36890 | 21.32 | 13.64 | 15.89 | 22.54 | 12.39 | 13.71 |
| *HMA3* | AT4G30120 | 0.31 | 0.7 | 0.78 | 1.05 | 0.48 | 0.81 |
| *FER1* | AT5G01600 | 102.43 | 78.55 | 117.63 | 108.02 | 14.4 | 22.31 |
| *FER3* | AT3G56090 | 56.85 | 47.51 | 57.41 | 35.61 | 19.2 | 22.4 |
| *FIT* | AT2G28160 | 6.45 | 7.58 | 2.97 | 13.17 | 9.68 | 2.53 |
| *bHLH38* | AT3G56970 | 0.01 | 0.01 | 3.21 | 14.94 | 0.05 | 41.64 |
| *bHLH100* | AT2G41240 | 0.01 | 0.11 | 5.5 | 26.57 | 0.01 | 72.05 |
| *bHLH39* | AT3G56980 | 2 | 3.17 | 5.6 | 19.06 | 84.61 | 39.79 |
| *bHLH101* | AT5G04150 | 0.01 | 0.01 | 2.76 | 11.09 | 24.21 | 29.23 |
| *NAS4* | AT1G56430 | 1.04 | 2.08 | 3.49 | 3.27 | 51.26 | 25.69 |
| *FRO3* | AT1G23020 | 20.23 | 27.23 | 17.27 | 45.19 | 251.49 | 130.05 |
| *OPT3* | AT4G16370 | 9.49 | 11.61 | 12.01 | 32.76 | 357.36 | 125.6 |
| *BTS* | AT3G18290 | 18.84 | 21.57 | 16.63 | 21.52 | 65.71 | 37.04 |
| *PYE* | AT3G47640 | 17.1 | 9.21 | 17.91 | 14.06 | 48.23 | 34.51 |
| *IMA1* | AT1G47400 | 0.11 | 0.32 | 0.11 | 4.68 | 29.38 | 13.37 |
| *IMA2* | AT1G47395 | 0.01 | 0.19 | 0.97 | 4.98 | 62.99 | 22.15 |
| *IMA3* | AT2G30766 | 0.09 | 0.25 | 0.18 | 0.98 | 29.08 | 10 |
| *IMA4* | AT1G07367 | 0.01 | 0.01 | 0.01 | 0.98 | 37.05 | 9.48 |
| *IMA6* | AT1G07373 | 0.01 | 0.01 | 0.15 | 3.02 | 56.06 | 31.12 |

The FPKM value of each gene is shown.

**Supplemental Table S2.** Expression of Fe-deficiency responsive genes in shoots.

| ID | AGI | +Fe | | | −Fe | | |
|---|---|---|---|---|---|---|---|
| | | WT | *bhlh4x-1* | *fit-2* | WT | *bhlh4x-1* | *fit-2* |
| *CYP82C4* | AT4G31940 | 0.01 | 0.01 | 0.01 | 0.44 | 0.01 | 0.01 |
| *S8H* | AT3G12900 | 0.04 | 0.09 | 0.05 | 0.31 | 0.01 | 0.01 |
| *NAS2* | AT5G56080 | 0.01 | 0.1 | 0.15 | 0.74 | 0.01 | 0.04 |
| *IREG2* | AT5G03570 | 0.01 | 0.04 | 0.01 | 0.26 | 0.18 | 0.04 |
| *MTP3* | AT3G58810 | 0.71 | 0.65 | 0.72 | 1.26 | 0.94 | 0.91 |
| *IRT1* | AT4G19690 | 0.08 | 0.1 | 0.53 | 4.44 | 1.36 | 0.81 |
| *FER1* | AT5G01600 | 244.37 | 277.83 | 132.8 | 111.15 | 64.23 | 61.82 |
| *FER3* | AT3G56090 | 37.98 | 39.44 | 28.57 | 14.52 | 7.35 | 9.98 |
| *FIT* | AT2G28160 | 0.18 | 0.12 | 0.13 | 0.81 | 0.13 | 0.24 |
| *bHLH38* | AT3G56970 | 0.12 | 0.18 | 4.05 | 21.17 | 0.01 | 67.99 |
| *bHLH100* | AT2G41240 | 0.39 | 0.01 | 17.33 | 54.5 | 0.08 | 240.68 |
| *bHLH39* | AT3G56980 | 0.05 | 0.26 | 5 | 11.12 | 68.91 | 51.24 |
| *bHLH101* | AT5G04150 | 0.49 | 0.17 | 4.79 | 7.19 | 19.14 | 46.08 |
| *PYE* | AT3G47640 | 7.05 | 14.21 | 17.29 | 43.44 | 87.15 | 60.37 |
| *BTS* | AT3G18290 | 25.89 | 21.94 | 39.36 | 40.51 | 152.18 | 131.17 |
| *MYB10* | AT3G12820 | 0.6 | 1.36 | 1.25 | 2.28 | 5.76 | 3.63 |
| *OPT3* | AT4G16370 | 56.63 | 45.75 | 87.35 | 76.61 | 271.78 | 184.46 |
| *FRO2* | AT1G01580 | 0.8 | 0.65 | 1 | 3.06 | 19.48 | 7.6 |
| *FRO3* | AT1G23020 | 38.68 | 61.7 | 64.13 | 141.93 | 354.8 | 209.18 |
| *NAS4* | AT1G56430 | 4.92 | 2.8 | 9.65 | 8.31 | 20.74 | 16.91 |
| *IMA1* | AT1G47400 | 0.01 | 0.3 | 2.56 | 14.72 | 59.1 | 33.88 |
| *IMA2* | AT1G47395 | 0.6 | 0.63 | 7.65 | 29.39 | 125.86 | 73.53 |
| *IMA3* | AT2G30766 | 1.02 | 2.03 | 20.89 | 56.79 | 144.25 | 112.99 |
| *IMA4* | AT1G07367 | 0.01 | 0.69 | 3.01 | 13.65 | 110.11 | 36.48 |
| *IMA6* | AT1G07373 | 0.01 | 0.4 | 3.62 | 13.26 | 141.96 | 51.88 |

The FPKM value of each gene is shown.

**Supplemental Table S3.** Primers used in this paper.

| Primer | Sequence (5'-3') |
|---|---|
| For qRT-PCR | |
| qFIT-F | CCAACACCTGTCGATGACCT |
| qFIT-R | TTCACCACCGGCTCTAACAC |
| qIRT1-F | GCCCCGCAAATGATGTTACC |
| qIRT1-R | TCCAATGACCACCGAGTGAA |
| qFRO2-F | ATCGAAAGTCGCCACACCAT |
| qFRO2-R | GAGCCACAAACATCGCCAAG |
| qNPTII-F | CATCATGGCTGATGCAATGC |
| qNPTII-R | CCCTGATGCTCTTCGTCCAG |
| qGFP-F | GCTGAAGGGCATCGACTTCA |
| qGFP-R | CCTTGATGCCGTTCTTCTGC |
| qACT2-F | TGTGCCAATCTACGAGGGTTT |
| qACT2-R | TTTCCCGCTCTGCTGTTGT |
| For amplification of full-length CDS | |
| FIT-F | ATGGAAGGAAGAGTCAACGC |
| FIT-R | TCAAGTAAATGACTTGATGAATTCA |
| bHLH38-F | ATGTGTGCATTAGTCCCTTCAT |
| bHLH38-R | CTAGTTAAACGAGTTTTCACATTTCT |
| bHLH39-F | ATGTGTGCATTAGTACCTCCATTG |
| bHLH39-R | TCATATATATGAGTTTCCACATTCC |
| bHLH100-F | ATGTGTGCACTTGTCCCTCC |
| bHLH100-R | TCATGTAAACGAGTGTCCACATT |
| bHLH101-F | ATGGAGTATCCATGGCTGCAG |
| bHLH101-R | TTATGATTGGCGTAATCCCAAG |
| For identification of bhlh4x-1 fit-2 | |
| fit-A | CCTTTTCGCGGTATCAATCCT |
| fit-B | TGCAGAACCGGATTTGACTCA |
| bhlh38-A | AGACACAAATGGGATCAAGTTG |
| bhlh38-B | AAGGGTTAACTCGGTGTTCTTC |
| b39-check-A | GCCATCAACGGGAGAGTACG |
| b39-check-B | AGCTTCTTCTGCAAGAAAGGAAA |
| bhlh100-A | TTGTGGTAGAAAAATGTGATTGC |
| bhlh100-B | TCAGTTTATGTTACTTGGGACCG |
| bhlh101-A | TTTCCTACTTCATCCCATCAAAG |
| bhlh101-B | TATGATTGGCGTAATCCCAAG |
